# Inter- and Intrahemispheric Sources of Vestibular Signals to V1

**DOI:** 10.1101/2024.11.18.624137

**Authors:** Guy Bouvier, Alessandro Sanzeni, Elizabeth Hamada, Nicolas Brunel, Massimo Scanziani

**Author notes:** The authors contributed equally to this work (Guy Bouvier, Alessandro Sanzeni).

## Abstract

Head movements are sensed by the vestibular organs. Unlike classical senses, signals from vestibular organs are not conveyed to a dedicated cortical area but are broadcast throughout the cortex. Surprisingly, the routes taken by vestibular signals to reach the cortex are still largely uncharted. Here we show that the primary visual cortex (V1) receives real-time head movement signals — direction, velocity, and acceleration — from the ipsilateral pulvinar and contralateral visual cortex. The ipsilateral pulvinar provides the main head movement signal, with a bias toward contraversive movements (e.g. clockwise movements in left V1). Conversely, the contralateral visual cortex provides head movement signals during ipsiversive movements. Crucially, head movement variables encoded in V1 are already encoded in the pulvinar, suggesting that those variables are computed subcortically. Thus, the convergence of inter- and intrahemispheric signals endows V1 with a rich representation of the animal’s head movements.

## Introduction

Many of the sensory organs through which we perceive the world are located in our head, for example, the eyes. To accurately represent our surroundings, sensory systems in the brain must combine their primary source of sensory information, e.g. visual signals, with information about head movements in space^1,2^. The vestibular organs, located in the inner ear, provide this information by transforming head movement into neural signals. Unlike other senses, however, these head movement signals are not processed by a dedicated cortical area, but are instead broadcast throughout the brain ^3–9^. Previous studies in rodents have demonstrated that neurons in primary sensory areas such as the primary visual cortex (V1) robustly respond to head movements, even in the absence of visual stimuli^6–8^. These responses depend on vestibular organs and dynamically track the time course of head movements, demonstrating specificity for aspects such as direction and velocity^7,8^. In contrast to our thorough understanding of the origin and processing of visual signals in V1, our understanding of head movement signals in this structure is still rudimentary. Is there a laminar organization in the representation of head movement signals in V1 as there is for visual information? Are head movement variables, such as direction and speed, computed in V1 or inherited from upstream structures? And, crucially, what are these upstream structures that relay head movement information to V1?

Here we use the mouse as a model system to determine the dynamics of V1 activity in response to head movement and reveal that the pulvinar nucleus of the thalamus, which receives axonal projections from the deep cerebellar nuclei (DCN), represents the main source of head movement signals to V1. We show that head movement variables, like direction and speed, are more accurately represented in the deep than the superficial layers of V1 and that, rather than being computed *ex novo* in V1, these variables are inherited from the pulvinar. The pulvinar, however, provides V1 with head movement signals that are biased toward contraversive movements (e.g. clockwise movements in left V1). Unexpectedly, we show that the contralateral visual cortex (VC) also provides V1 with head movement signals which, in contrast to the pulvinar, are stronger during ipsiversive head movements, and thus counterbalance the pulvinar bias. These results show that V1’s rich representation of an animal’s head movement variables results from the integration of inter- and intrahemispheric signals.

## Results

### Encoding of Head Movements in V1

We recorded extracellular activity in the left V1 of head-fixed, awake mice in response to vestibular stimulation, delivered in the dark, by rotating the animal along the horizontal plane with a servo-controlled rotating table (Figure 1A, top). This protocol elicits responses in V1 that entirely depend on the vestibular organs^7,8^. Most V1 neurons (63%; 1490/2355 neurons; N = 37 mice) responded to clockwise (CW; i.e. contraversive relative to left V1: 49%; 1152/2355 neurons) and/or counterclockwise (CCW; i.e. ipsiversive relative to left V1: 47%; 1112/2355 neurons) rotations of the table, by either increasing or decreasing their firing rate (FR), as described previously (Methods for class assignment criteria and statistical tests throughout).

**Figure 1.**
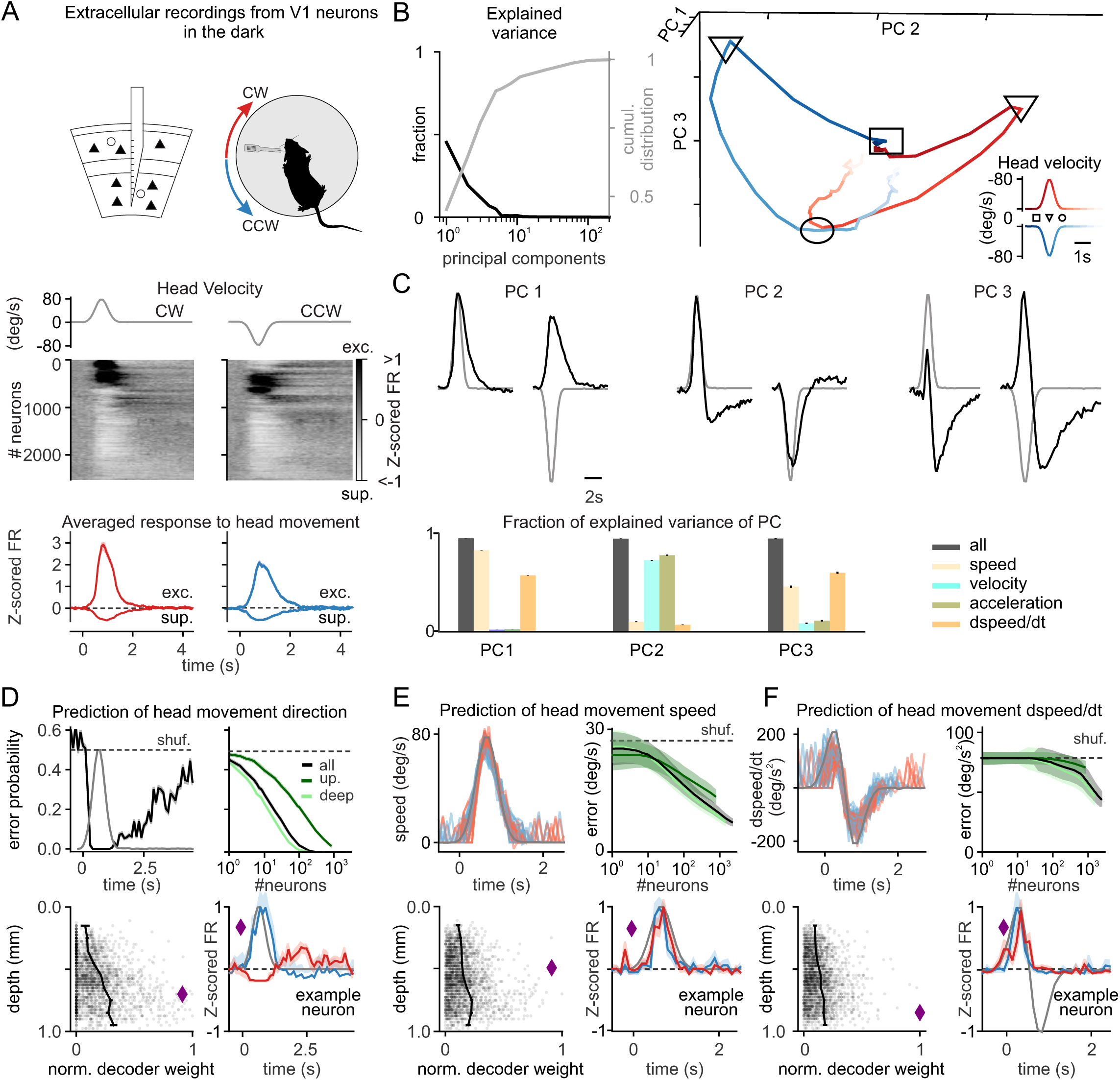
V1 encodes head movements in the absence of visual stimuli. (A) Top: experimental configuration. A linear probe spanning all cortical layers was inserted in the left V1 of a head-fixed, awake mouse to record the response to clockwise (CW) and counterclockwise (CCW) rotations of a table in the dark. Middle: UMAP sorting of the responses, represented by the averaged Z-score of the firing rate (Z-scored FR) across neurons. Neurons were ordered using UMAP with cross-validation. Bottom: Average Z-scored FR of neurons significantly excited (exc.) or suppressed (sup.) by head movement in CW (red) and CCW (blue) direction. The gray traces on top are the velocity profile. (B) Principal component (PC) analysis of V1 population activity during head movements. Left: fraction of explained variance (black) and its cumulative distribution (gray) as a function of increasing PC. Right: Three-dimensional representation of the first three PCs. Note the separation of CW (red) and CCW (blue) representations. Iinset: time is color coded, and square, triangle and circle symbols indicate beginning, peak, and end of the rotation, respectively. (C) Top: population dynamics along the first three PCs (black) superimposed on the velocity profile (gray). Bottom: variance explained in each PC by models including speed, velocity, and their time derivatives either all together (gray) or separately (color). (D) Representation of head movement direction in a trial. Top left: decoding error probability as a function of time. Top right: decoding error probability as a function of the number of neurons (computed at peak velocity). Bottom left: decoder weight of individual neurons (gray dots) and population average (black connected dots; bin = 100 μm) as a function of depth. Bottom right: example neuron (purple diamond) selected using the highest decoder weight for head movement direction, at the peak and 2 s after the peak velocity (red: CW; blue: CCW. (E) Prediction of head angular speed. Top left: decoding head angular speed as a function of time. Top right: error as a function of number of neurons (color indicates depth as in D). Bottom left: decoder weight of single neuron as a function of depth. Bottom right: example neuron (purple diamond) selected using highest decoder weight for head angular speed decoding (red: CW; blue: CCW). (F) Analogous to E, but for the time derivative of speed.

Head movements trigger compensatory eye movements in the opposite direction via the vestibulo-ocular reflex (VOR). Since eye movements are known to modulate V1 neuronal activity^10,11^, we tested whether V1 neurons can respond to head movements independently of eye movements. For this, we implemented a vestibular stimulation protocol that eliminates VOR, called VOR cancellation (Methods). Here, instead of rotating the animal in the dark, we display a vertical grating on a virtual drum that rotates along the horizontal plane together with the animal, thereby preventing compensatory eye movements (Figure S1). Even during VOR cancellation, 66% of V1 neurons responded to head movements (231/351 neurons; N = 5 mice; P = 0.18 compared to vestibular stimulation in the dark). Thus, the activity of V1 neurons is strongly modulated by head movements even in the absence of eye movements. All subsequent experiments were conducted in the dark.

Among left V1 neurons that responded to CW and/or CCW rotations, changes in FR were, on average, larger for CW than CCW rotations (|Z-scored FR| in CW and CCW rotations were: 1.16 ± 0.05 including 1152/1490 neurons and 0.85 ± 0.03 including 1112/1490 neurons, respectively; P = 4.5e^-3^). Furthermore, more than half of these neurons (56%; 834/1490 neurons) showed a significant direction preference, responding more strongly to head movements in one direction (Methods), as previously reported^7^. Among this population, neurons preferring CW rotations were slightly, yet significantly overrepresented (55%; 455/834 neurons prefer CW rotations; P = 3.5e^-3^). To determine whether the preference for CW rotations in left V1 represents hemispheric specialization, we recorded from V1 neurons in the right hemisphere. Neurons in right V1 showed a bias toward CCW head rotation, thus opposite to left V1. Specifically, among right V1 neurons showing significant direction preference, the proportion of neurons preferring CW rotations was significantly lower than in left V1 (45%; 48/106 neurons; P = 1.4e^-3^), while the proportion of neurons preferring CCW rotations was similar to that for CW rotations in left V1 (55%; 58/106 neurons; P = 0.4). These results reveal that in V1, neurons respond to head movements with a slight overrepresentation for contraversive rotations.

V1 neurons’ response to head movements exhibited diverse amplitudes, kinetics, and signs (Figure 1A), and the first five PCs captured 89% of the variance (Figure 1B; variance per PC ≥ 0.05; Methods). To explore the population dynamics, we visualized activity along the first three PCs (Figure 1B). Before head movement onset, activity was confined to a small neural space. As speed increased, distinct trajectories emerged for CW and CCW movements, consistent with the direction preference of V1 neurons described above. Interestingly, the population dynamics observed with decreasing speed did not overlap with that of increasing speed, and this differential representation persisted for several seconds after the head movement ceased (3.7 s; P<0.05; Figure 1B,C). These results suggest that V1 encodes information about head movements beyond movement direction and instantaneous speed, potentially including changes in speed over time and the history of previous movements.

To determine which aspects of head movement were encoded in each PC, we fitted their dynamics using a linear model of speed, angular velocity (CW and CCW rotations as positive and negative velocities), and their derivatives. This model captured the dynamics of all five PCs (Figure 1C; explained variance ≥ 0.72). Fitting each variable individually revealed that speed and its derivative explained 86% and 60% of PC1 variance, angular velocity and acceleration explained 75% and 80% of PC2 variance, and speed and its derivative explained 46% and 61% of PC3 variance. Thus, V1 dynamics reflect head angular velocity, speed, and their derivatives.

To test if head movement variables could be extracted on a trial-by-trial basis, we developed a decoding analysis. Logistic regression models predicted movement direction (CW or CCW) using neural activity in 100 ms bins (Figure 1D; Methods). With all recorded neurons (n = 2355), decoding error was at chance before movement onset but dropped below 0.05% within 300 ms of initiation, remaining above chance for 3.7 s after offset. Accuracy exceeded 99.5% with fewer than 600 neurons (Figure 1D), with deeper-layer neurons (below 500 µm) contributing more than superficial ones (above 500 µm). Notably, single high-weight neurons could predict direction with over 99.5% accuracy (Figure S2A). The speed of head movements was accurately decoded from V1 activity (error = 6.8 ± 0.8 deg/s with 2355 neurons in 100 ms bins), with deeper-layer neurons providing more information (Figure 1E). Unlike direction decoding, where high-weight neurons maintained their responses for seconds post-movement, the neurons decoding speed closely tracked instantaneous head speed, achieving similar accuracy with just ∼30 neurons (11.9 ± 2.1 deg/s; Figure S2A). Derivatives of speed, velocity, and acceleration were decoded with similar performance (Figures 1F, S2B-C). Decoders generalized across head movement profiles, performing consistently across phases of velocity or varying accelerations (Figure S2D,E).

Taken together, these results show that mouse V1, and especially its deeper layers, encodes a rich representation of head movement that can be accessed to simultaneously decode both present and past movements with high precision.

### Pulvinar origin of head movement signals to V1

What are the sources of the head movement signals that reach V1? To identify potential upstream candidate areas, we injected an anterograde transsynaptic tracer in the DCN, one of the main sources of vestibular signals to the brain. Injection of AAV2/1-hSyn-Cre in the DCN of a tdTomato reporter mouse labelled neurons throughout the thalamus, in agreement with previous work^12–14^. Among visual areas, only the pulvinar nucleus of the thalamus receives direct DCN projections (Figure 2A). In contrast, DCN injections labelled very few neurons in the dorsal lateral geniculate nucleus (dLGN), the other main thalamic relay to visual cortex, nor did they label neurons in V1 (pulvinar: 896.5 ± 97.8 per mm^3^; dLGN: 27.8 ± 5.7 per mm^3^, P = 6.6e^-17^; no labelled neurons in V1; N = 8 mice), consistent with the lack of direct projection from DCN to cortex^12–14^. We confirmed the specificity of the DCN projections to the pulvinar using a retrograde tracer. Injection of retrograde AAV-Cre in the pulvinar of tdTomato reporter mice labelled more neurons in the DCN than in the other main source of vestibular signals – the vestibular nuclei (DCN: 682.4 ± 82.7 per mm^3^; VN: 94.5 ± 27.0 per mm^3^; P = 2.7e^-11^; N = 3 mice; Figure S3A-C). Thus, pulvinar neurons receive direct projections from the DCN, making this thalamic nucleus a potential node for vestibular signals on their way to V1.

**Figure 2.**
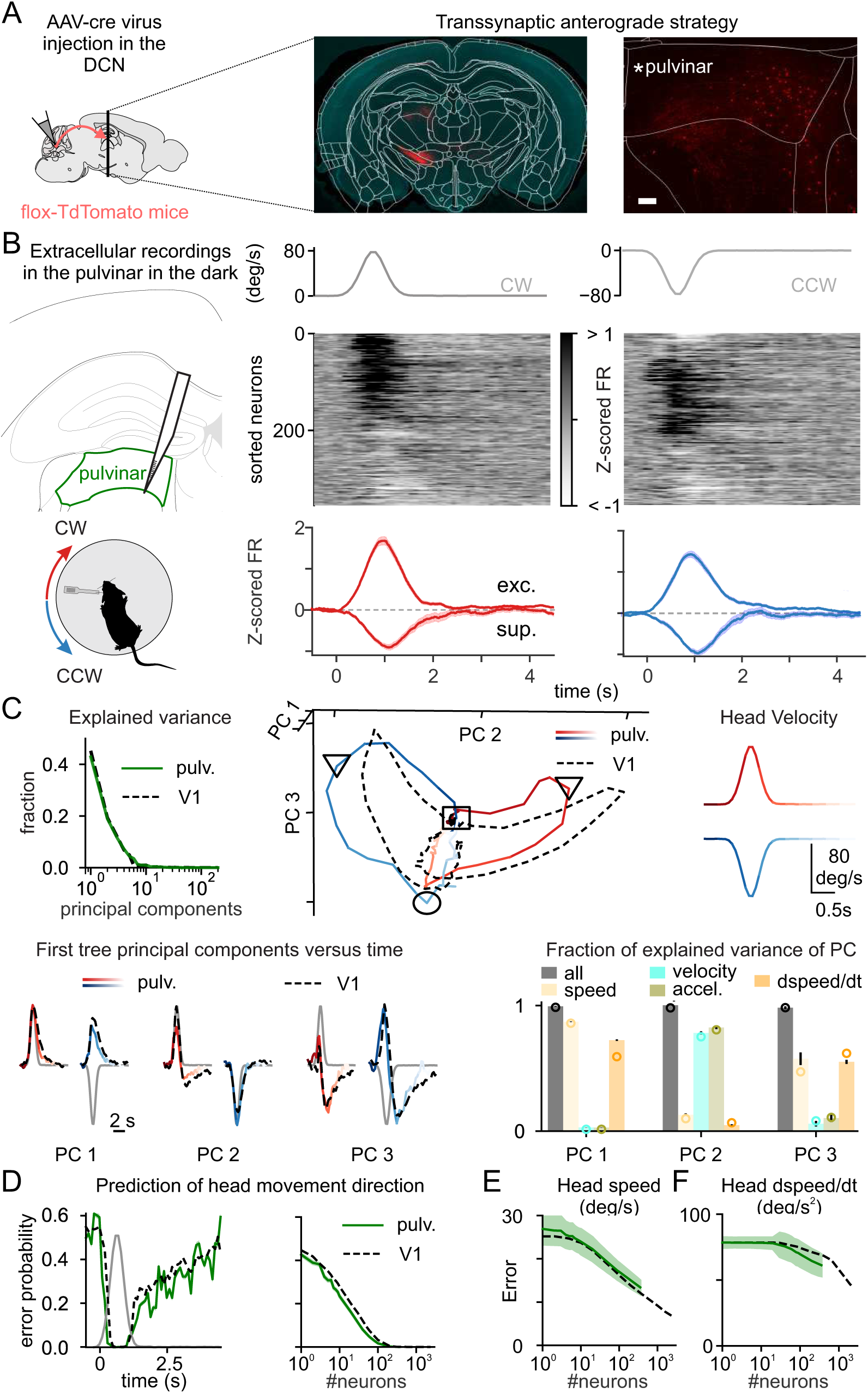
The pulvinar receives input from the deep cerebellar nuclei and encodes head movements like V1. (A) Experimental strategy. Left: Injection of Cre dependent transsynaptic anterograde virus (AAV2.1 hSyn-cre) in the deep cerebellar nuclei (DCN) in a flex-tdTomato reporter line; this approach labels the projections of the DCN and the post-synaptic neurons in red (Methods). Middle: Photomicrograph of coronal sections at −1.98 mm from Bregma illustrating transynaptically labelled neurons. Right: zoom on the left pulvinar thalamus. Scale bar = 100 μm. (B) Experimental configuration. Left: extracellular linear probe in the left pulvinar of a head-fixed, awake mouse records the response to clockwise (CW, red) and counterclockwise (CCW, blue) rotations of the table in the dark. Right: UMAP sorting of the averaged Z-score of the firing rate (Z-scored FR) responses across neurons (neurons were ordered using UMAP with cross-validation), and Z-scored FR of pulvinar neurons that significantly respond to CW (red) and CCW (blue) head movements. The gray traces on top are the velocity profile. (C) Comparison of principal component (PC) analysis in pulvinar and V1. Top left: Fraction of explained variance in the pulvinar (green) and in V1 (black, as described in Fig. 1B). Top right: Three-dimensional representation of the first three PCs for pulvinar activity (time is color coded: CW (red) and CCW (blue) rotations; square, triangle and circle symbols indicate beginning, peak, and end of the rotation, respectively) and V1 activity (dashed lines). Bottom: Variance explained in each PC by models including speed, velocity, and their time derivative either all together (gray) or separately (color). Circles are the fraction of explained variance in V1 (from Fig. 1C). (D) Decoding error probability of head movement direction. Left: as a function of time in a trial. Right: as a function of the number of neurons (right, computed at peak velocity) in pulvinar (green) and V1 (black). (E) Decoding error of head angular speed as a function of the number of neurons in pulvinar (green) and V1 (black). (F) As in E, but for head angular derivative of speed in time.

If the pulvinar is a source of head movement signals to the visual cortex, it must respond to head movements. Thus, we recorded neuronal activity from the left pulvinar of head-fixed, awake mice in response to vestibular stimulation delivered in the dark by rotating the animal along the horizontal plane (Figure 2B), as we did for V1 recordings (see above). Similar to V1, the firing rate of most pulvinar neurons (73%; 258/355 neurons; N = 11 mice) was modulated by CW (58%; 207/355 neurons) and/or CCW (51%; 182/355 neurons) rotations of the table and a large fraction of these neurons (45%; 160/355 neurons) exhibited a significant direction preference (Figure 2B). Furthermore, the response of left pulvinar was also biased toward CW head movements (average Z-scored FR for CW 1.36 ± 0.10 and CCW 1.04 ± 0.08; P = 3.7e^-3^). In fact, the pulvinar showed a larger fraction of neurons preferring CW rotations than V1 (62% including 100/160 neurons in the pulvinar versus 55% including 455/834 neurons in V1; P = 3.7e^-2^). Finally, principal component analysis of pulvinar activity during head movement revealed a remarkable similarity with that observed in V1 (Figure 2C). Specifically, the activity along the first three PCs (Figure 2C, top right and bottom) closely matched those observed in V1. Moreover, the first five PCs were explained by head movement-related variables in an analogous way to V1 (Figure 2C). The similarity between head movement representation properties in the pulvinar and V1 were equally striking when quantified with a decoding analysis. Decoding performances of trial head movement direction as a function of time (Figure 2D), as well as of speed (Figure 2E), velocity (Figure S3E), and their time derivatives (derivative of speed: Figure 2F; acceleration: Figure 3F), closely resemble what we observed in V1.

**Figure 3.**
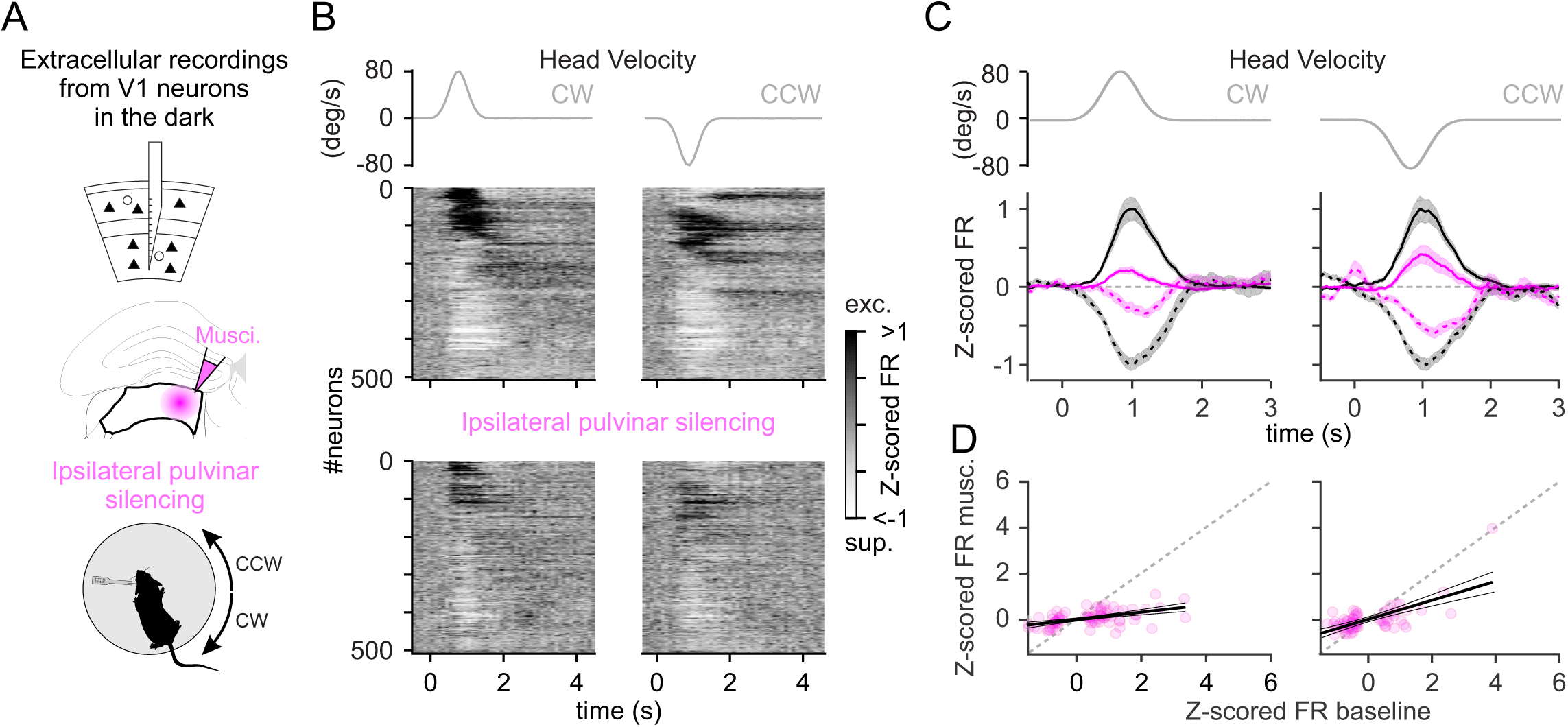
V1’s response to head movements depends on the ipsilateral pulvinar. (A) Experimental configuration. Top: linear probe spanned all cortical layers, before and after ipsilateral pulvinar silencing by injecting muscimol-BODIPY. Bottom: extracellular linear probe in the left V1 of a head-fixed, awake mouse records the response to clockwise (CW) and counterclockwise (CCW) rotations of the table in the dark. (B) UMAP sorting of the averaged Z-score of the firing rate (Z-scored FR) of V1 neurons. Left: response to CW head movements. Right: response to CCW head movements. Middle: in control condition before pulvinar silencing. Bottom: after pulvinar silencing. Neurons were ordered so that nearby ones had similar Z-scored FR over time (UMAP with cross-validation; see Methods). The gray traces on top are the velocity profile. (C) Peak normalized Z-scored FR of V1 neurons. Left: response to CW head movements. Right: response to CCW head movements. In control condition (dark traces) and after ipsilateral pulvinar silencing (magenta traces). Note that for C and D, only neurons significantly modulated during head movements under control condition contribute to each average depending on whether they were excited (solid lines) or suppressed (dashed lines) by the rotation. All traces are normalized by the peak of the Z-scored FR under control condition. The gray traces on top are the velocity profile. (D) The peak Z-scored FR. Left: response to CW head movements. Right: response to CCW head movement. Control condition (x-axis) versus pulvinar silencing (y-axis). Magenta circles represent neurons that significantly respond to head movements under control conditions.

Taken together, these results demonstrate that head movements modulate the activity of a large fraction of pulvinar neurons to generate a rich representation that matches the one observed in V1, albeit with a stronger bias toward contraversive movements. Thus, the pulvinar is a potential source of vestibular signals upstream of V1.

To determine whether the pulvinar contributes to the representation of head movements in V1, we pharmacologically silenced this thalamic area while simultaneously recording V1 activity in response to vestibular stimulation. The stereotactic injection of the GABAergic agonist muscimol into the left pulvinar resulted in a slight decrease in the basal activity of V1 neurons (control: FR = 3.77 ± 0.29 Hz (average ± sem); pulvinar silencing: FR = 3.44 ± 0.26 Hz; P = 1.0e^-3^; n = 358 neurons, N = 4 mice). Yet, it caused a strong reduction of their response to head movements (CW: 72 ± 5% decrease of Z-scored FR; P = 1.5e^-8^; CCW: 54 ± 6% decrease of Z-scored FR; P = 1.7e^-5^; Figure 3). This effect was more pronounced for CW rotations (CW versus CCW: P = 6.0e^-3^, Figure 3), consistent with the biased representation of CW head movements in the pulvinar (see above).

Taken together, these results indicate that the pulvinar is a main source of vestibular signal to V1 with a bias toward contraversive head movements.

### Contralateral visual cortex contribution of head movement signals to V1

The fact that silencing the left pulvinar reduces responses in left V1 to CW more than to CCW rotations suggests that left V1 receives CCW head movement signals from an additional source. Because responses in right V1 are biased toward CCW rotations, and V1 hemispheres are connected via transcallosal projections^15,16^, we tested the potential contribution for right V1 to CCW head movement signals in left V1.

For this, we optogenetically silenced the right visual cortex while recording from left V1 (Figure 4A). Silencing was achieved by photo-activating inhibitory neurons expressing Channelrhodopsin2 (ChR2; VGat-ChR2-EYFP mouse line) with an LED placed on top of the right visual cortex, as described previously^17–19^. Silencing trials were alternated with control ones without LED illumination. We prevented LED-evoked visual responses (i.e., LED light hitting the retina) to contaminate our V1 recordings by performing experiments in mice previously blinded by intraocular TTX injections in both eyes (Methods). LED illumination led to a slight decrease in the average basal activity of left V1 neurons (control FR = 2.78 ± 0.20 Hz (average ± sem); contralateral VC silencing FR = 2.70 ± 0.19Hz; P = 1.8e^-10^; n = 408 neurons; N = 7 mice; 23% (95/408) of the neurons were suppressed and 19% (75/408) excited). Strikingly, silencing the right visual cortex selectively reduced responses to CCW head rotations in left V1 (33.2 ± 5.7% decrease of Z-scored FR; P = 8.0e^-10^) leaving responses to CW rotations unaffected (CW: 11.6 ± 5.7% decrease of Z-scored FR; P = 0.83; CW versus CCW: P = 1.0e^-4^; Figure 4A-E). Thus, the right visual cortex is a source of CCW head movement signals to left V1.

**Figure 4.**
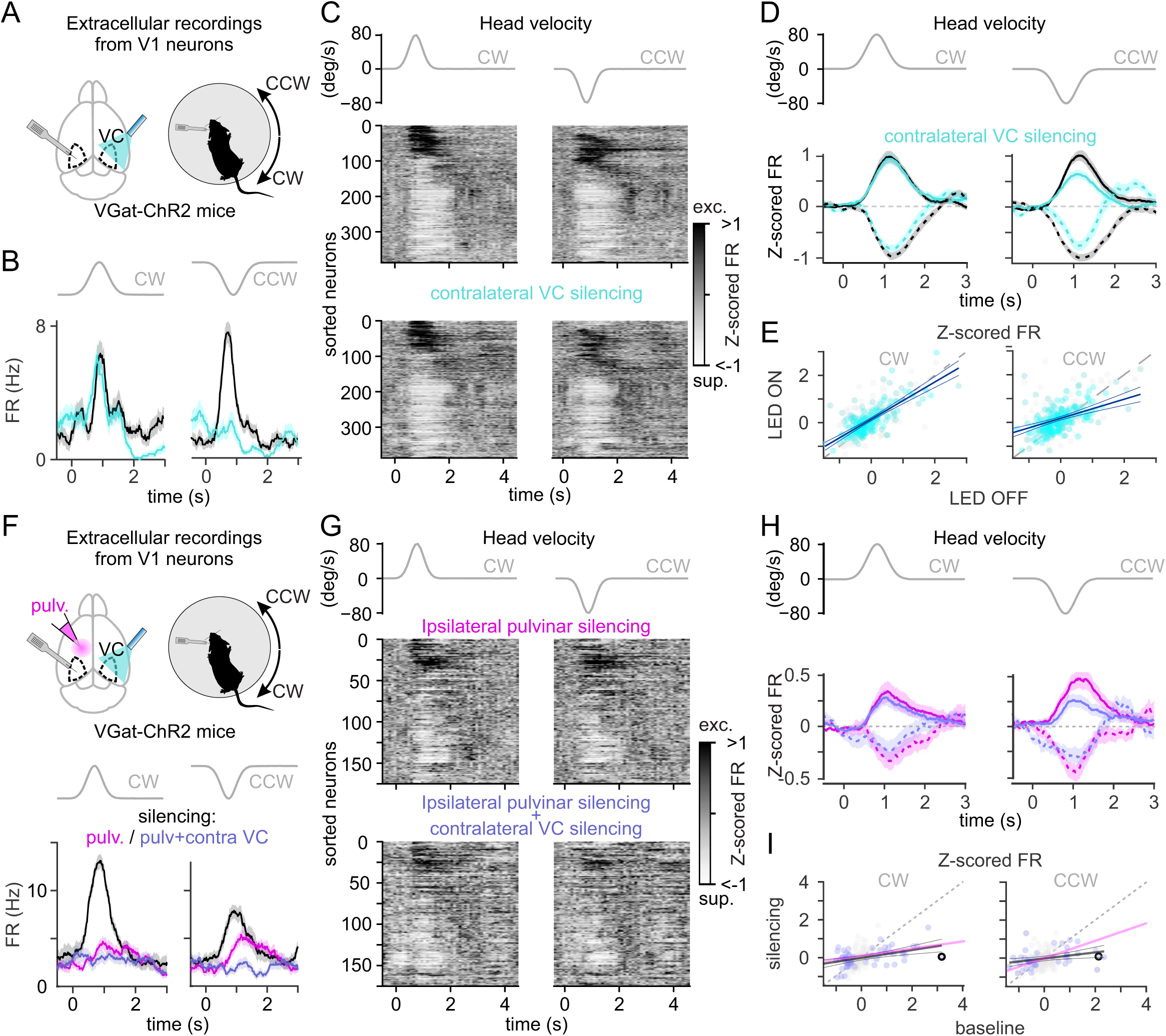
Ipsilateral pulvinar and contralateral visual cortex are sources of complementary vestibular signals to V1. (A) Experimental configuration. Left: extracellular linear probe positioned in the left primary visual cortex (V1) while optogenetically silencing the contralateral visual cortex (VC) using LED illumination through a thinned skull on a VGat-ChR2-EFP transgenic mouse. Right: extracellular linear probe in the left V1 of a head-fixed, awake mouse records the response to clockwise (CW) and counterclockwise (CCW) rotations of the table while optogenetically silencing the contralateral visual cortex. (B) Effect of silencing contralateral VC on an example neuron recorded in left V1. Z-score of the firing rate (Z-scored FR). Left: responses to CW head movements. Right: responses to CCW head movements. Control condition (dark traces) and during contralateral VC silencing (cyan traces). The gray traces on top are the velocity profile. (C) UMAP sorting of the averaged Z-scored FR of V1 neurons. Left: responses to CW head movements. Right: response to CCW head movements. Middle: in control condition. Bottom: during contralateral VC silencing. The gray traces on top are the velocity profile. (D) Peak normalized Z-scored FR of V1 neurons. Left: responses to CW head movements. Right: responses to CCW head movements. Control condition (dark traces) and during contralateral VC silencing (cyan traces). (E) Relation between Z-scored FR in control versus contralateral VC silencing conditions. Same neurons as in D. (F) Experimental configuration. Left: extracellular linear probe positioned in the left V1 before and after pharmacological silencing of the ipsilateral pulvinar (muscimol-BODIPY), while the contralateral optogenetically VC (same as A) is silenced on alternate trials. Right: extracellular linear probe in the left V1 of a head-fixed, awake mouse records the response to CW and CCW rotations of the table while silencing the ipsilateral pulvinar and the contralateral visual cortex. Bottom: Effect of silencing contralateral VC and ipsilateral pulvinar on an example neuron. Z-scored FR of a V1 neuron. Left: response to CW head movements. Right: response to CCW head movements. In control condition (dark traces), during ipsilateral pulvinar silencing (magenta traces), and during simultaneous ipsilateral pulvinar and contralateral VC silencing (purple traces). (G) UMAP sorting of the averaged Z-scored FR of V1 neurons. Left: in responses to CW head movements. Right: in response to CCW head movements. Top: after ipsilateral pulvinar silencing only. Bottom: after both ipsilateral pulvinar and contralateral VC silencing. The gray traces on top are the velocity profile. (H) Peak normalized Z-scored FR of V1 neurons. All traces are normalized by the peak Z-scored FR observed under control condition. Left: response to CW head movements. Right: response to CCW head movements. Following ipsilateral pulvinar silencing (magenta traces) and during contralateral VC and pulvinar silencing (purple trace). The gray traces on top are the velocity profile. Note that the control condition is not represented. (I) Relation between Z-scored FR in control versus simultaneous contralateral VC silencing and ipsilateral pulvinar silencing conditions. Bold circles report the example neuron in F. Note that for D and H, only neurons significantly modulated during head movements in the control condition contribute to each average depending on whether they were excited (solid line) or suppressed (dashed line) by the rotation. All traces are normalized by the peak Z-scored FR observed in control condition. The gray traces on top are the velocity profile.

If the ipsilateral pulvinar and the contralateral cortex independently contribute to head movement responses in V1, CCW responses remaining in left V1 after pulvinar silencing should be further reduced by silencing the right visual cortex. We tested this possibility by combining pharmacological silencing of the pulvinar with the optogenetic silencing of the visual cortex. Consistent with this hypothesis, optogenetic silencing of the right visual cortex performed following the pharmacological silencing of the left pulvinar selectively reduced the remaining responses to CCW rotations in left V1 (CW: 14.3 ± 14.5% decrease of Z-scored FR; P = 0.61; CCW: 25.0 ± 11.2% decrease of Z-scored FR; P = 0.03; n = 175 neurons; N = 5 mice; Figure 4F-H).

Taken together, these results indicate that the ipsilateral pulvinar is the main contributor to V1 responses to head movements, with a bias toward contraversive rotations while the contralateral visual cortex contributes to responses to ipsiversive rotations. Thus, the ipsilateral pulvinar and, to a lesser extent, the contralateral visual cortex are two independent sources of vestibular signals to V1, each preferentially contributing to responses in opposite directions along the horizontal plane.

## Discussion

Primary sensory areas in the cerebral cortex process modality-specific sensory information originating from peripheral receptors. Recent studies, however, have challenged this strict, modality-specific view^7,10,20–24^. A striking example is that V1 responds robustly to head rotations via vestibular organ activation^7,8^, even in complete darkness. Thus, V1 is a primary sensory cortical area that not only receives ascending input from the eyes and responds to visual stimuli, but also responds to vestibular stimuli. This observation has opened several fundamental questions, relative to the anatomical pathways taken by these vestibular signals to reach V1, the nature of the representation of head movement variables in V1 and the extent to which the representation of these variables is inherited from upstream structures. Here, we demonstrate that V1 accurately encodes head movement variables (direction, speed, velocity, and their time derivatives) especially in deeper layers, and that it receives head movement signals through two main pathways: the ipsilateral pulvinar nucleus of the thalamus and the contralateral visual cortex. The ipsilateral pulvinar provides the predominant head movement signal, exhibiting a bias toward contraversive rotations (e.g., clockwise rotations relative to left V1). In contrast, the contralateral visual cortex contributes head movement signals during ipsiversive rotations. Importantly, we found that head movement variables in V1 are already represented in the pulvinar, suggesting that V1 inherits these variables rather than computing them *ex novo*. These results demonstrate that the integration of intra- and interhemispheric signals endows V1 with a rich and accurate representation of head movements.

Sensory stimuli often trigger behavioral responses, which, in turn, can lead to a cortex-wide modulation of neuronal activity^21,25^. Consequently, some activity in primary cortical areas in response to stimuli of distinct modalities may be erroneously interpreted as multimodal responses. For example, a substantial portion of V1 neurons’ response to auditory stimuli can be explained by facial movements triggered by the sound, and are thus not true auditory V1 responses^23,24^. In our experiments, head movements also triggered a behavioral response, namely compensatory eye movements through the activation of the VOR^26^. Given that the kinetics and amplitudes of both head and eye movements are tightly linked during VOR, V1 responses to head movement could, in theory, be explained by eye rather than head movements. This possibility would be consistent with the fact that the VOR, like the response of V1 to head movements, depends on the vestibular organ^7,8^. To address this possibility, we used a well-established protocol to eliminate eye movements during head movement, called VOR cancellation^27^. Using this protocol, we still recorded strong responses of V1 neurons to head movements, demonstrating that these responses arise from the head movement itself rather than from concurrent compensatory eye movements.

The thalamus receives the majority of its vestibular input from the contralateral vestibular and cerebellar nuclei,^12–14^ and these nuclei are primarily biased toward ipsiversive movements. Consistent with this organization, we observed a contraversive movement bias in the pulvinar (i.e., clockwise for the left pulvinar). However, this contraversive bias of the pulvinar almost disappears in V1. We discovered that this rebalancing of motion specificity is due to input from the contralateral visual cortex. While the cortico-callosal projections in the binocular V1 have been well characterized^28^, their role in monocular V1 is still unclear^15,16^. Here, we show that this interhemispheric communication to monocular V1 provides a head movement signal with a direction preference bias opposite to that obtained from the ipsilateral pulvinar. Whether this rebalancing is critical for a complete (ipsi- and contralateral) picture of motion, or if the segregation and recombination of motion signals serves a specific computational purpose remains to be seen.

Several pathways connect the vestibular system to the primate’s thalamus^29^, and accordingly, the pulvinar of these animals responds to various vestibular stimuli^30^. Furthermore, the pulvinar receives projections from motor areas, including the motor cortex and deep cerebellar nuclei^12,13,31^, and projects broadly across the visual cortex, conveying not only visual information but also other non-visual inputs to V1^10,32^. Interestingly, inactivating the pulvinar abolishes the saccadic modulation observed in V1 neurons^10^, highlighting the pulvinar’s role in integrating both visual and non-visual information. Our results demonstrate that, in mice, the pulvinar serves as the primary source of head movement signals to V1. The convergence of visual information in V1 with head and eye movement signals conveyed through the pulvinar suggests a fundamental mechanism for maintaining stable perception during active behavior. Indeed, the pulvinar has recently emerged as critical for distinguishing self-generated from external visual motion^33^, and its role in predictive coding^34^ may reflect a broader strategy where the brain uses movement-related signals to anticipate sensory consequences. Future work will reveal how the pulvinar’s diverse functions arise from its unique position between motor and sensory systems, and how this integration shapes our perception of a stable visual world during self-motion.

In summary, the intra- and interhemispheric vestibular signals to V1 described here may impact cortical visual processing by providing a head movement context for incoming visual inputs. Understanding the origin and processing of these signals could offer critical insights into how the early visual system integrates visual input with contextual information related to motion, thereby enhancing our understanding of sensory processing during action.

## METHODS

### EXPERIMENTAL MODEL AND SUBJECT DETAILS

#### Mice

All experimental procedures were conducted in accordance with the regulations of the Institutional Animal Care and Use Committee (IACUC, AN179056) of the University of California, San Francisco.

All mice were housed on a reversed cycle (light/dark cycle 12/12 h) with free access to food. Data were collected from male or female C57BL/6J mice or from heterozygous mice kept on a C57BL/6J background with the following genotype: VGat–ChR2–EYFP (Jackson Labs #014548). V1 recordings in darkness included 1,502 units from 30 C57BL/6J mice, obtained from our previous study under identical experimental conditions^7^. At the start of the experiments, all mice were between 2 and 7 months old.

### METHODS DETAILS

#### Viruses

The following adeno-associated viruses (AAV) were used: AAV1-hSyn-Cre-WPRE-hGH (final titer: 1.8×10^13^ genome copies/ml, Univ. of Pennsylvania Viral Vector Core) and AAV1-retro-hSyn-Cre-eBFP (final titer: 5×10^12^ genome copies/ml, Univ. of Pennsylvania Viral Vector Core).

#### Surgical procedures

##### Viral Injections

Mice were anesthetized with 2% isoflurane and placed in a stereotactic apparatus (Kopf). Core body temperature was monitored with a rectal probe and maintained constant at 37°C with a heating pad (FHC). A thin layer of lubricant ointment (Rugby Laboratories) was applied to the eye, the head was shaved and disinfected with povidone iodine, and 2% lidocaine solution was administered subcutaneously at the incision site. A craniotomy (approx. 300 µm in diameter) was performed with a micro-burr (Gesswein) mounted on a dental drill (Foredom). Viral suspensions were loaded in beveled glass capillaries (tip diameter: 15-30 µm) and injected with a micropump (UMP-3, WPI) at a rate of 30-40 nl/min into the parenchyma. The coordinates of the injection sites and the volumes of the injected viral suspension are detailed below. The pipette was removed from the brain 15 min after the completion of the injection, the head plate was attached just after the virus injection, and 0.1 mg/kg buprenorphine was administered subcutaneously as a postoperative analgesic. For anterograde transsynaptic strategy^35^, the virus was injected in the right DCN (AP: −6.1 mm; ML: 2.0 mm; depth: 2.25 mm; volume = 100-250 nl). For retrograde strategy in the pulvinar, the virus was injected in the left rostro-medial pulvinar (AP: −1.9 mm; ML: 1 mm; depth: 2.4 mm; volume = 30-50 nl).

##### Head Plate Implantation for Head-fixed Recordings

Mice were implanted with a T-shaped head-bar at least 2.5 weeks before the day of the recording. Mice were anesthetized with 2% isoflurane, the scalp was removed, the skull was disinfected with alcohol and povidone iodine, and scored with bone scraper. The edge of the skin was glued to the skull and the metal head-bar was sterilized and mounted using dental cement (Ortho-Jet powder; Lang Dental) mixed with black paint (iron oxide), or Relyx Unicem2 automix (3M ESPE). The head-bar was stereotactically mounted with the help of an inclinometer (Digi-Key electronics 551-1002-1-ND). The inclinometer allowed us to adjust the angle of the head bar in relation to the sagittal and medio-lateral axes of the head. Following the bar implantation, black dental cement was used to build a recording well surrounding the recording site. The surface of the skull above the left visual cortex was not covered with dental cement but was coated with a thin layer of transparent cyanoacrylate glue. Mice were injected subcutaneously with 0.1 mg/kg buprenorphine and checked daily after the head-bar surgery. For at least 4 days before recording, mice were habituated to head fixation within the recording setup.

##### Craniotomy for Electrophysiological Recordings

On the day before recording, mice were anesthetized with 2% isoflurane and the skull above the recording sites was drilled off. The dura was not removed, and the exposed brain was kept moist with artificial cerebrospinal fluid (ACSF; 140 mM NaCl, 5 mM KCl, 10 mM D-glucose, 10 mM HEPES, 2 mM CaCl2, 2 mM MgSO_4_, pH 7.4). V1 recordings were performed at approximately 2.6 mm lateral to the sagittal suture and 0.6 mm anterior to the lambdoid suture.

#### Electrophysiology

Extracellular recordings were performed using the following silicon probes Neuronexus: A1×32-5mm-25-177-A32; A1×32-Edge5mm-20-177-A32; A2×32-5mm-25-177-A64, 1×64-Poly2-6mm-23 s-160 or Cambridge Neurotech: ASSY-77 H2 (Acute 64 channel H2 probe, 2 shanks @250 µm, 8 mm length), ASSY-77 H5 (Acute 64 channel H5 probe, 1 shank, 9 mm length). The recording electrodes were controlled with Luigs & Neumann micromanipulators and stained with DiI or DiO lipophilic dyes (Thermo Fisher) for post hoc identification of the electrode track. We recorded the signals at 30 kHz using an INTAN system (RHD2000 USB Interface Board, INTAN Technologies).

#### Head-fixed Rotations

To control the velocity and amplitude of head movements, we fixed the head of awake mice in the center of a servo-controlled platform enabling the rotation of the animal along the horizontal plane (50 degrees rotation; 80 deg/s peak velocity, see Fig. 1A; unless stated otherwise). Mice were head-fixed, their bodies restrained in a tube, and we pseudo-randomly alternated clockwise (CW) with counterclockwise (CCW) rotations. The platform was attached to a gearbox 15:1 (VTR010-015-RM-71 VTR, Thomson) that increased the torque of a servo motor (AKM53L-ANC2C-00 KEC0432 AC Servomotor 1.83 kW, Kollmorgen). The motor was tuned using a servo drive (AKDB013206-NBAN-0000 servo drive, Kollmorgen) and controlled in velocity mode using analog waveforms computed in Labview.

#### Monitoring eye movements by video-oculography

The movement of the right eye was monitored through a high-speed infrared (IR) camera (Imperx Bobcat, B0620). The camera captured the reflection of the eye on an IR mirror (transparent to visible light, Edmund Optics #64–471) under the control of custom Labview software and a frame grabber (National Instrument PCIe-1427). The pupil was identified online or post hoc by thresholding pixel values and its profile was fitted with an ellipse to determine the center. The eye position was measured by computing the distance between the pupil center and the corneal reflection of a reference IR LED placed along the optical axis of the camera. To calibrate the measurement of the eye position, the camera and the reference IR LED were moved along a circumference centered on the image of the eye by ± 10 degrees^36^.

#### Vestibulo-ocular reflex paradigms

To assess vestibulo-ocular reflex (VOR) compensation and cancellation, well-habituated mice were head-fixed on a rotating platform surrounded by a visual virtual stimulus drum^36^. We presented visual stimuli (0.1 cpd) moving synchronously with the turntable (20 deg peak velocity, 1.8 s, 15 deg) during VOR cancellation and static during VOR compensation trials. Eye movements were tracked as described above. During VOR cancellation trials, the platform and visual drum were rotated using a gaussian velocity waveform in the same direction and at matching velocities. This condition required mice to suppress their VOR to maintain a stable gaze on the moving visual stimulus. Eye position data were analyzed offline using custom MATLAB scripts to calculate gain (ratio of eye and head velocity). VOR cancellation performance was quantified as the absence of reflexive eye movement, while expecting a reflexive eye movement in the opposite direction during VOR compensation. Rapid eye movements were excluded from our analysis.

#### Pharmacology

Intraocular injection of tetrodotoxin (TTX; 40 µM) was performed 2 hours prior to recording, under isoflurane anesthesia. A typical procedure lasted less than five minutes. TTX was injected in both eyes for all the experiments performed on VGat-ChR2-EYFP mice. Immediately prior to the injection, a drop of proparacaine hydrochloride ophthalmic solution was applied to the eye as a local anaesthetic (Bausch + Lomb, 0.5%). TTX solution was injected intravitreally using a beveled glass micropipette (tip diameter ∼50 µm) on a micro injector (Nanoject II, Drummond) mounted on a manual manipulator. 1 µl was injected in each eye, at the speed of 46 nl/s. The animals were head-fixed for recording following a 2-hour recovery period in their home cage.

Silencing of the pulvinar was performed by injecting 30-40 nl of 5 mM muscimol-BODIPY at the speed of 80-150 nl/min, using a beveled glass pipette (tip diameter ∼ 20-40 µm) on a micro injector UMP3 with a Micro4 controller (World Precision Instruments). The injector was mounted on a micromanipulator (Luigs & Neumann) for stereotactic injection. After the recording, brains were fixed in 4% PFA in PBS overnight at 4°C for histological analysis of BODIPY on the next day.

To verify the absence of visual responses following intraocular TTX injection, we used a full-field luminance change from 0 cd.m^−2^ to 100 cd.m^−2^ lasting 1 s.

#### Optogenetic silencing of contralateral visual cortex

Cortical silencing was achieved by expressing channelrhodopsin-2 (ChR2) in inhibitory neurons, a technique previously validated^17–19^. We utilized the VGat–ChR2–EYFP mouse line for optogenetic silencing of the contralateral visual cortex. For photostimulation of ChR2-expressing cortical inhibitory neurons, we positioned a 470-nm blue fiber-coupled LED (1 mm diameter, Doric Lenses) approximately 5-10 mm above a thinned skull area on the right hemisphere of the visual cortex. To limit illumination to the tissue under the cranial window, we covered adjacent areas with black dental cement. To prevent inadvertent retinal stimulation from blue light, we induced temporary blindness by injecting both eyes with TTX (see above). The LED fiber delivered a total light power of 8-15 mW. We alternated trials between head rotation alone and head rotation combined with LED illumination. The LED was activated for 4 s, centered on the peak velocity of the head rotation.

#### Histology

For anatomical analysis, mice were transcardially perfused with phosphate buffered saline (PBS) and then with 4% paraformaldehyde (PFA) in PBS. Brains were extracted from the skulls, post-fixed in 4% PFA overnight at 4°C, and subsequently cut with a vibratome to 80-100 µm thick sequential coronal sections. Slices were collected and mounted in ProLong Gold (Life Technologies) or Vectashield mounting medium containing DAPI (Vector Laboratories H1500). Bright-field and fluorescence images were acquired using an Olympus MVX10 MacroView microscope. For quantifying the number of somata in visual thalamus, deep cerebellar nuclei, and vestibular nuclei (see sections: *Viruses* and *Viral Injections*), neuronal density was counted for each brain slice (visual thalamus: n = 123 slices; deep cerebellar and vestibular nuclei: n = 40 slices) and then averaged. Ipsilateral projections to the injection site, for both DCN and pulvinar tracing experiments were negligible and not considered. The Paxinos brain atlas was used as a reference to delineate these regions.

### QUANTIFICATION AND STATISTICAL ANALYSIS

#### Data Analysis

##### Unit isolation

Automated spike sorting was carried out using KiloSort and KiloSort2 (https://github.com/cortex-lab/Kilosort) by manual curation of the units using Phy and Phy2 (https://github.com/cortex-lab/phy). Single units were identified, and all the following analysis was carried out via MATLAB (MathWorks), but for principal component and decoding analysis (Python 3). The quality of the isolated units was assessed using refractory period violations and stability of amplitude.

##### Class assignment criteria

To classify the units that are significantly modulated by head movement, we compared its neuronal activity before and during head rotation. The baseline spike rate was calculated on individual trials by averaging the spike rate over a window of 580 or 1000 ms recorded when the platform was stationary before the rotation. The spike rate in response to the rotation of the platform was calculated on the same trials by averaging the spike rate over a window of 580 ms centered around the peak of the rotation velocity profile. Wilcoxon signed-rank tests were then applied to determine if a unit was significantly modulated (P < 0.05) by the rotation of the platform. Directional preference of individual unit was quantified by comparing their firing rate in response to CW and CCW rotations and using non-parametric Wilcoxon signed-rank tests, with statistical significance set at P < 0.05. This analysis enabled identification of neurons exhibiting preferential responses to specific rotation directions. For each recorded unit, we computed the mean firing rate within a 580 ms temporal window centered on the peak angular velocity of platform rotation. To compare the Z-score of the firing rate collected from 2 populations of mice, we performed Wilcoxon rank sum tests. When reporting averaged absolute Z-score of the firing rate, only neurons significantly modulated by head movement in baseline condition were included. Wilcoxon signed rank tests were performed to compare values obtained in the same recording and only neurons significantly modulated in control condition for a given direction of rotation were considered (i.e., comparing before and during thalamus or contralateral visual cortex silencing).

##### Cortical depth estimation

Cortical depth from pia estimated by using electrophysiological landmarks across layers as described previously^7,37^. Briefly, the Multi-unit (MUA) spectral power (500 Hz to 5 kHz) distribution along the probe track allowed us to locate layer 5a. This approach allowed us to normalize the cortical depth from the pia across mice.

##### Statistics

Statistical analyses were done using MATLAB and Python 3. No statistical tests were used to predetermine sample size, but our sample sizes are similar to those generally employed in the field. All data are presented as mean ± standard error of the mean (SEM), unless otherwise noted. The stated P values are the results of the non-parametric Wilcoxon rank sum test to compare values between different mice or recordings, and the non-parametric Wilcoxon signed rank test to compare values from the same recording in different experimental conditions. The difference of fraction of neurons modulated across brain areas was accessed using the bootstrap hypothesis test. Specifically, we resampled neurons with replacement from our dataset of Z-score of the firing rate responses to clockwise (CW) and counterclockwise (CCW) rotations (N = 10000 iterations). For each bootstrap sample, we computed response amplitudes and firing rate changes, then fitted Gaussian functions to the cumulative distribution functions to calculate p-values, determining the statistical significance of observed differences. For the anatomical tracing analysis, the non-parametric Wilcoxon rank test was employed to compare the values between slices from the VN and the DCN, as well as between slices from the dLGN and the pulvinar. Experiments and analyses were not blinded.

##### Cross validated neuron sorting and principal component analysis

For recordings in V1 and pulvinar in control conditions, trials were divided in two equally populated sets (called in what follows training and test sets); a trial averaged response was computed for each neuron separately in the two sets. Neurons were ordered using the UMAP algorithm applied to the training set of the control conditions^38^. This ordering was then applied to the test set to obtain the UMAP plots shown throughout the text.

Cross-validated principal component analysis activity^21^ was used to estimate the fraction of variance explained by each principal component. In brief, we computed the trial averaged response of neurons in each realization of the training and test sets. The training set was used to derive the principal components, while the test set was used to measure the variance explained in each component. The procedure was repeated ten times with different random realizations of the training and test sets; means and standard errors over realizations are shown in Figure 1B, 1C, and 2C. As shown by Stringer et al.^21^, this method measures the reliable variance of stimulus-related dimensions, excluding trial-to-trial variability from unrelated cognitive and/or behavioural variables or noise. For recordings in V1 and the pulvinar, the first 3 (pulvinar: 5) PCs accounted for 76% and 74% (pulvinar: 87% and 85%) of the variance, respectively.

##### Relationship between principal components and movement

To quantify the relation between movement and neural activity along each principal component (PC), we defined a predictor y(t) given by

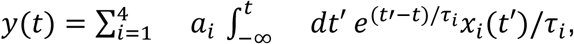

where the index *i* runs over the movement related variables investigated (speed, velocity, dspeed/dt, acceleration), *x*_*i*_(t) is the temporal profile of the *i*-th variable, and each of these variables are convolved with an exponential kernel of amplitude *a*_*i*_ and time constant *τ*_*i*_. This choice was motivated by the fact that single neuron and population dynamics showed long lasting responses. For each PC, we fitted the corresponding values of *a*_*i*_ and *τ*_*i*_ by minimizing the square difference between the measured population dynamics and *y*(*t*). Analogously with what described for cross validated PCA, fits were performed on a training set, validation was measured on a test set, the procedure was repeated 10 times with random realizations of training and test sets, and the mean and standard error of the predictor performance (measured with R^2^) were evaluated averaging over realizations. In V1 and the pulvinar, the model captures a large amount of variance in the first 4 PCs (97.2 ± 0.2, 96.2 ± 0.7, 93.7 ± 2.0, and 88.0 ± 1.3 in V1; and 95.9 ± 0.9, 93.3 ± 1.1, 85.5 ± 4.0, and 80.5 ± 4.8 in the pulvinar), but then much less in other PCs (55.7 ± 4.0 in V1 and 18.1 ± 17.8 in the pulvinar).

Trial to trial variability in the single neuron responses affected our estimates of firing rates and led to different dynamics along each PC in the training and the test sets. These fluctuations are due to the finite number of trials in the experiments; hence they could not be captured by our “kernel” model described above, but can strongly influence our estimate of its performance. To account for this phenomenon in our quantification of the predictor performance, we defined a “null model”, which used the dynamics observed in the training set as predictor for the dynamics along each PC in the test set. This null model quantifies the reliability of our estimate of firing rates. It was used as a reference to evaluate the performance of the “kernel” model. Specifically, in each PC we computed the proportion of the variance in the test set explained by the null model (measured with *R*^2^). A value *R*^2^ close to 1 indicates that the estimate of the firing rate of neurons was reliable across training and test sets; a value close to zero or negative, on the other hand, indicates that our estimates of firing rates were mainly determined by trial to trial fluctuations. We found that the null model in V1 and the pulvinar had a positive *R*^2^ only for the first 5 and 6 components, respectively. For PCs with positive *R*^2^, we computed the ratio between *R*^2^ given by the kernel model and the null model; this ratio, which we called the fraction of explainable variance captured, is shown in the bar plots of Figures 1C and 2C.

To estimate the importance of the *i*-th variable, we repeated the procedure described above, setting *a*_*j*_ =0 for all *j* ≠ *i*. The analysis in V1 revealed that the first PC was mostly explained by speed and its derivative (Kernel time constants 0.219 ± 0.004 s and 5.0 ± 0.1 s); the second PC was mostly explained by velocity and acceleration (kernel time constants 8.56 ± 0.05 ms and 4.8 ± 0.1 s); while the third PC was mostly related to speed and its time derivative (kernel time constants 2.62 ± 0.03 s and 0.674 ± 0.008 s). Similar results were obtained in the pulvinar.

##### Decoding analysis

Decoding of head movement related information from neural activity was performed by training decoders on spike counts in bins of 100 ms. Decoders were trained on 80% of the bins and tested on the remaining 20%. Figures in the manuscript only show decoder predictions on test bins. To evaluate performance of decoders, training and testing were repeated 100 times, randomly shuffling which trials were used for training and for testing. Decoding performance with shuffled labels were evaluated using the same procedure, but shuffling the association between spikes in a bin and the corresponding animal head movement. Numerical analyses were performed using the python library scikit-learn.

Logistic regression models (solver ‘lbfgs’ and ‘l2’ regularization)^39^ were used to decode head movement direction. To quantify the history dependent movement response, we trained separate decoders for each bin to predict if the corresponding trial was CW or CCW. Training was performed assigning sample weight to each movement value (CW or CCW) that corresponded to its frequency in the dataset. The regularization parameter C of each decoder was determined by maximizing the cross validated performance. Generalization performance across movement profiles was measured using a unique logistic regression model for all the time bins along the trial.

Velocity and acceleration decoding were performed using a nonlinear decoder constructed combining a logistic regression model predicting instantaneous movement direction (three categories: no movement, CW, CCW) and two distinct ridge regression models (corresponding to bins with CW and CCW movements) predicting instantaneous movement magnitude. Unlike those used to characterize history dependent effects, the logistic regression models used here were unique for all the time bins along the trial. The ‘l2’ regularization parameter of the ridge regression model was optimized to maximize cross validation performance. Training was performed assigning sample weight to each data point; these were computed dividing possible head movement values in 30 equally spaced bins and measuring the frequency of each movement bin in the dataset. We found that this nonlinear decoder outperformed a simpler linear decoder, obtained with a single ridge regression model; this result was likely due to the fact that, unlike the linear decoder, the nonlinear decoder was able to exploit neurons with symmetric CCW and CW response.

To evaluate performance of decoders as a function of the number of neurons, we systematically measured decoding performance as a function of the number of neurons the decoder had access to. For a fixed number of neurons, this was implemented by randomly picking which neurons were used in the decoding and repeating the procedure 1000 times. Mean and standard error of the decoding performance over random realizations of the training and test set are shown in Figures 1D-F, S2B-D, 2D-F, and S3E,F. In the top-left panel of Figures 1E, 1F, S2B, S2C, and S2E, as well as in all of Figure S2D, the individual colored lines represent independent predictions, each generated from random realizations of the training and test sets.

## Acknowledgments

We thank Pooja Saraf, Madan Mukundan, Brandon Wong, Qui Ying Wu, and Leo Ruan for technical assistance, the former and current members of the Scanziani lab for discussions, and the members of the SensoMotion lab for critical reading of the manuscript.

## Funding

National Institutes of Health grants U19NS107613 (MS), R01EY025668 (MS), U01NS108683 (AS and NB) and Howard Hughes Medical Institute (MS).

### Author contributions

G.B. and M.S. designed experiments. G.B., A.S., N.B., and M.S. wrote the manuscript. G.B. performed all experiments, except that E.H. processed the tissues for tracing experiments. G.B. and A.S. analyzed the data. A.S. performed all the principal components and decoding analysis.

---

**Figure S1.**
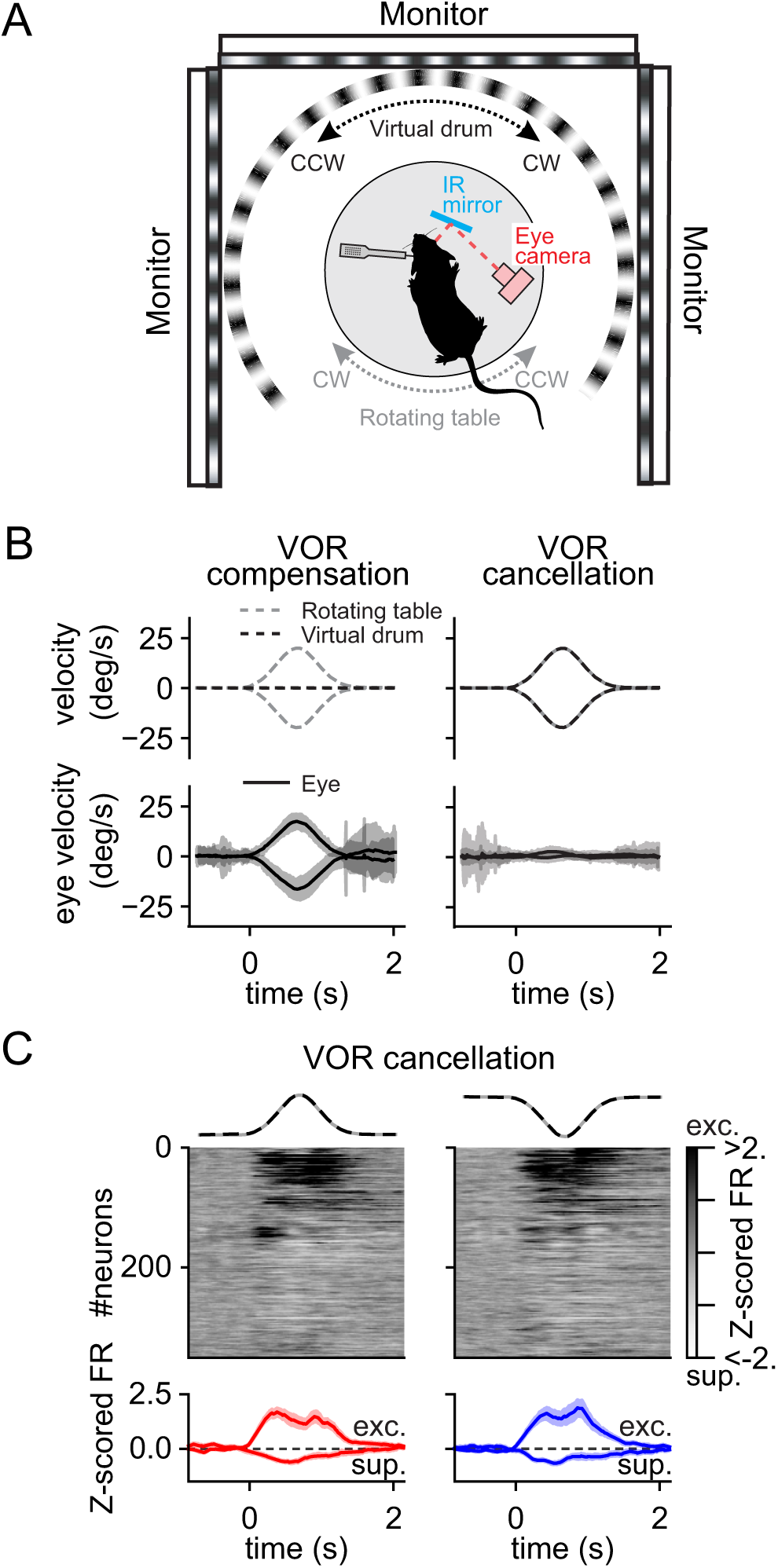
V1 neurons respond to head movements with or without compensatory eye movements, related to. **Figure 1**. (A) Experimental configuration. Extracellular linear probe in the left V1 of a head-fixed, awake mouse records the response to clockwise (CW) and counterclockwise (CCW) rotations of the table (gray dotted line) surrounded by a virtual drum made of a pattern of vertical light and dark stripes (black dotted line). A camera monitors the right eye through an infrared (IR) mirror. To minimize the number of resetting saccades, we use a lower peak velocity (20 deg/s) as compared to that used in all other experiments (Methods). (B) Vestibulo-ocular reflex (VOR) under two conditions: compensation (left) and cancellation (right). Top left: VOR compensation is triggered by table rotations (dotted gray line) in front of a static virtual drum (dotted black line). Bottom left: VOR compensation is characterized by eye movements in the opposite direction but same angular amplitude of the rotating table. Top right: VOR cancellation is triggered by simultaneous rotation of the virtual drum and the rotating table in the same direction. Bottom right: VOR cancellation is characterized by the absence of compensatory eye movements. (C) V1 neurons responses during VOR cancellation. Top: UMAP sorting of the average Z-scored FR responses across neurons performing CW (left) and CCW (right) head movements during VOR cancellation. Bottom: Z-scored average of the firing rate for the neurons that are significantly excited (exc.) and suppressed (sup.) by CW (red) and CCW head movement (blue). The gray and black dotted traces on top are the velocity profiles of the table and of the virtual drum, respectively.

**Figure S2.**
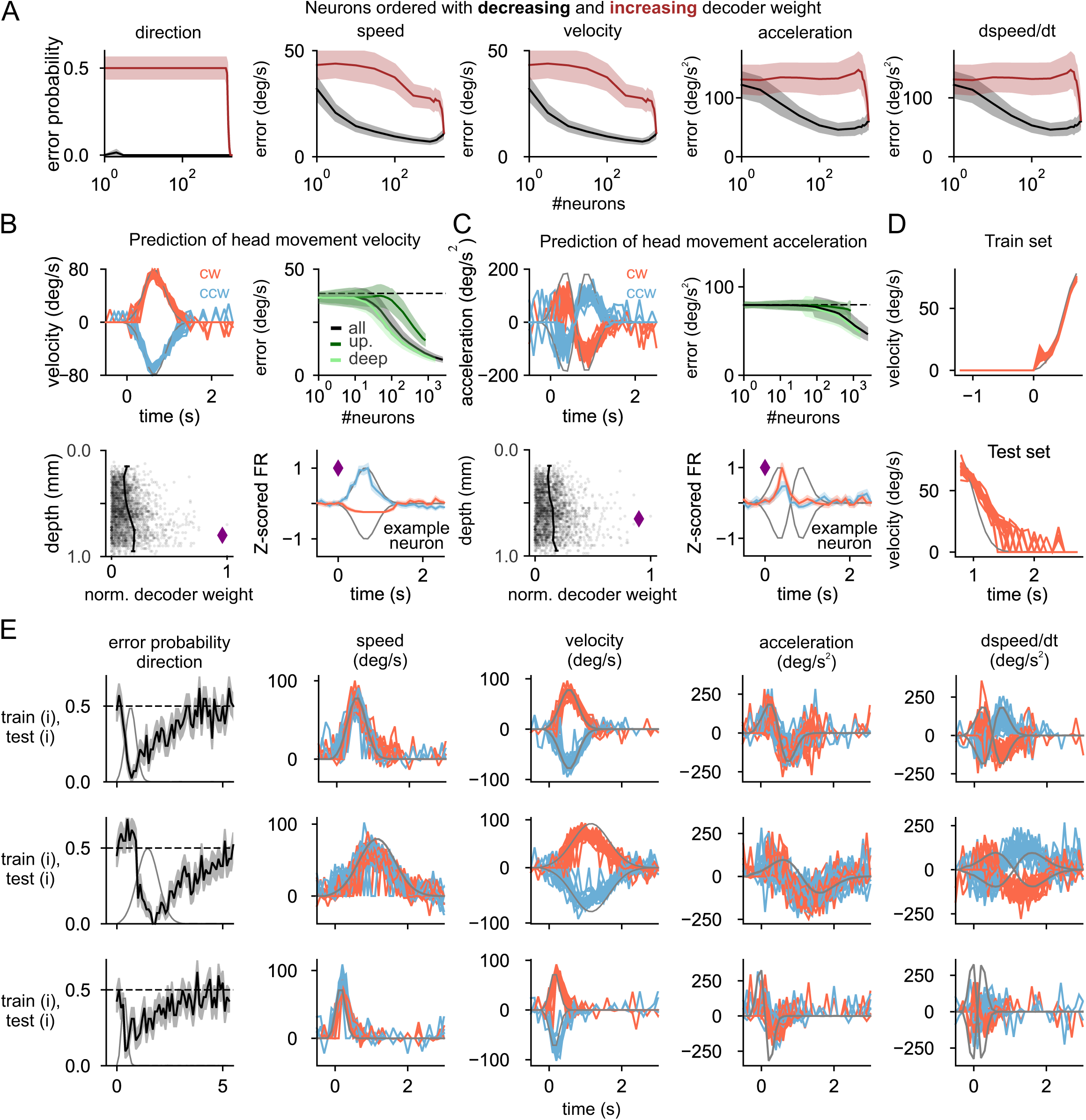
Prediction of head angular velocity and acceleration and generalization of head movement information to different head angular rotation profiles, related to Figure 1. (A) Decoding error of head angular rotations as a function of the number of neurons. Neurons were ranked according to decoding weight in decreasing (black) or increasing (red) order. The decoding weight was obtained with all neurons available to the decoder (i.e. those shown in Fig. 1 D-F of the main text). The errors shown for a given N in the plots are those obtained from a decoder which has access to the first N neurons in the ranking order. (B) Top left: prediction of head angular velocity. Top right: error as a function of the number of neurons. Bottom left: decoder weight of single neuron as a function of depth. Bottom right: example neuron (purple diamond) selected using highest decoder weight for head angular speed decoding. In the top-left panel, as in the following panels, colored lines represent independent predictions, each derived from random realizations of the training and test sets. In the top-right panel, light green and dark green indicate deep and superficial layers, respectively. (C) As in B, but for time derivative of velocity (acceleration). (D) Top: prediction of head angular velocity of a decoder trained on the rising phase of the head movement profile. Bottom: prediction of head angular velocity of a decoder tested on the decaying phase of the head movement profile. (E) Top line: Instantaneous head movement direction, speed, velocity, acceleration, and derivative of speed predicted by a decoder trained on profiles with 80 deg/s peak velocity. Middle line: tested on slower velocity profiles with the same peak velocity. Bottom line: tested on faster velocity profiles with the same peak velocity. Example neurons in panels A-C are marked with purple diamonds. In A-C and E, blue and red traces correspond to quantities measured during CCW and CW rotations, respectively.

**Figure S3.**
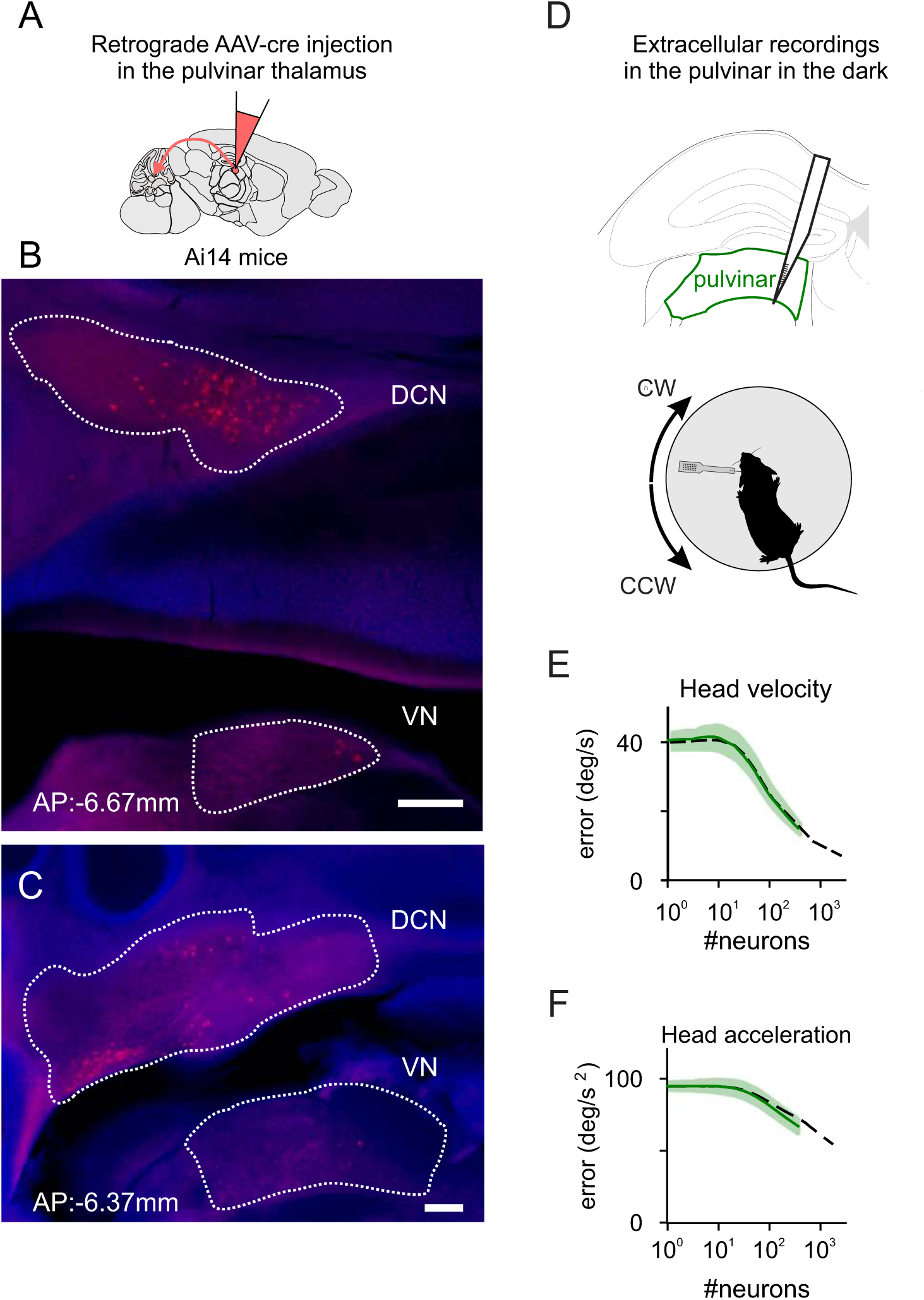
The pulvinar receives projections from the deep cerebellar nuclei and encodes head angular velocity and acceleration, related to. **Figure 2**. (A) Experimental strategy. Injection of Cre dependent retrograde virus (rAAV2.1 hSyn-cre) in the rostro-medial pulvinar in a flex-tdTomato reporter line; this approach labels the neurons that project to the pulvinar. (B, C) Photomicrograph of coronal sections at −6.67mm (B) and −6.37mm (C) from Bregma illustrating retrogradely labelled neurons in the deep cerebellar and vestibular nuclei, respectively. DAPI is in blue and tdTomato is red. Scale bar: 200 μm. (D) Experimental configuration. Top: Extracellular linear probe spanned the ipsilateral pulvinar in the dorso-ventral axis. Bottom: extracellular linear probe in the pulvinar (green) of a head-fixed, awake mouse records the response to CW (red) and CCW (blue) rotations of the table in the dark. (E) Decoding error of head angular velocity as a function of number of neurons in pulvinar (green) and V1 (black). (F) As in E, but for head angular acceleration.

## Notes

### Competing Interest Statement

The authors have declared no competing interest.

### Summary of Updates

No significant difference between this version and the previous version.

